# Molecular hydrogen can minimize negative effects of heat stress on the hard coral genus *Acropora*

**DOI:** 10.1101/2024.08.02.606431

**Authors:** Malte Ostendarp, Mareike de Breuyn, Yusuf C. El-Khaled, Neus Garcias-Bonet, Susana Carvalho, Raquel S. Peixoto, Christian Wild

## Abstract

Coral reefs are increasingly threatened by mass bleaching events due to global ocean warming. Novel management strategies are urgently needed to support coral survival until global efforts can mitigate ocean warming. Given the strong antioxidant, anti-inflammatory and anti-apoptotic properties of molecular hydrogen, our study explores its potential to alleviate the negative effects of heat stress on corals. We investigated the ecophysiological responses of two common hard corals (*Acropora* spp. and *Pocillopora verruco*sa) from the Central Red Sea under ambient (26 °C) and elevated seawater temperatures (32 °C), with and without hydrogen addition (∼ 150 µM H_2_) over 48 h. Our results showed that at 32 °C without hydrogen addition, *P. verrucosa* exhibited high temperature tolerance, whereas *Acropora* spp. showed significant reductions in photosynthetic efficiency and maximum electron transport rate compared to the ambient condition (26 °C). The addition of hydrogen at 32 °C increased the maximum electron transport rate of *Acropora* spp. by 28 %, maintaining it at levels compared to those at 26 ° C. This suggests that molecular hydrogen benefits specific coral species under heat stress in the short-term. This study provides the foundation for future long-term and in-situ studies, potentially guiding the development of new management strategies aiming at enhancing coral resilience to ocean warming.

## Introduction

Coral reefs are among the most biodiverse and productive ecosystems worldwide [1]. They provide humankind with a diverse range of essential ecosystem services [1], on which millions of livelihoods depend [2]. However, coral reefs are currently in serious decline [3, 4], as a result of many different local and global anthropogenic and natural threats, including local pollution, overfishing and global warming [5, 6, 7, 8, 9].

Although many of these threats are nowadays identified and partially addressed, climate change continues at a rapid pace with minimal progress made towards mitigation [10, 11]. In association with the onset of El Niño in 2023 [12], new ocean temperature records are emerging [13, 14] greatly increasing the probability and frequency of mass coral bleaching and mortality events worldwide [10, 15, 16, 17]. Direct actions and strategies are therefore urgently needed to further prevent the rapid degradation of these highly valuable ecosystems thereby gaining time for coral reefs until climate change becomes manageable by global efforts [18].

Shifts in physical and biogeochemical conditions are widely recognized to predominantly induce coral bleaching referring to the loss of the coral’s endosymbiotic algae Symbiodiniaceae often representing the main energy source of the coral host [19, 20]. Novel approaches could therefore aim to target cascades associated with coral bleaching, particularly those initiated by heat stress, offering a promising approach for intervention. Accordingly, several studies indicate that oxidative damage is one driver behind coral bleaching as proposed by the “Oxidative Theory” [21, 22].

In this context, molecular hydrogen might offer an effective treatment since it was discovered as a great preventive and therapeutic agent in human medicine attributed to hydrogen’s strong antioxidant, anti-inflammatory and anti-apoptotic characteristics [23, 24, 25, 26, 27, 28]. As a strong antioxidant, molecular hydrogen selectively reduces hydroxyl radicals and peroxynitrite belonging to the most harmful ROS/RNS [23]. Concurrently, other ROS/RNS with a potentially important physiological role were not affected [23, 27, 28]. Since molecular hydrogen is also able to rapidly diffuse across membranes, it can successfully penetrate cell organelles [23] and scavenge ROS/RNS inside the cytoplasm and mitochondria. In addition, molecular hydrogen is proposed to inhibit the generation of ROS/RNS by preventing leakage of electrons from the electron transport chain [28], regulate gene expressions [27], increase overall antioxidant capacity [29, 30] and upregulate the heat shock response [31].

Based on these previously observed effects of molecular hydrogen in mammals including humans and rats [23, 24, 25, 26, 27, 28], here we provide first insights into the potential effects of molecular hydrogen on the coral holobiont answering following research question: How does molecular hydrogen affect the ecophysiology of corals under heat stress? Considering its antioxidant, anti-inflammatory and anti-apoptotic properties [23, 24, 25, 26, 27, 28], we hypothesize that molecular hydrogen minimizes negative effects of heat stress.

To address the research question and associated hypothesis, we experimentally examined the short-term effect (48 h) of hydrogen-enriched seawater under ambient temperature and heat stress on the two common and widely distributed hard corals *Acropora* spp. and *P. verrucosa* from the Central Red Sea.

## Materials and Methods

### Coral collection and experimental preparation

To assess the short-term effect of hydrogen addition on the coral holobiont, two coral species *Acropora* spp. and *P. verrucosa* commonly found in the Red Sea were selected. Coral fragments were sampled by SCUBA diving near the “Coral Probiotics Village” at Al Fahal Reef (22°18’19.1’’N, 38°57’55.0’’E), located about 20 km off the King Abdullah University of Science and Technology, Thuwal, in the Saudi Arabian coast of the Red Sea. Permits were obtained from the Institutional Biosafety and Bioethics Committee of the King Abdullah University of Science and Technology (IBEC protocol number 22IBEC003_v4).

A total of six colonies per species were marked at a depth of approximately 10 m to allow the identification and resampling of the same colonies during the experiment duration. On each sampling day, two coral fragments per colony measuring about 3 - 5 cm in length were collected using pliers along with 90 L of seawater in water containers. Due to laboratory space limitations, it was not possible to run all incubations simultaneously. Therefore, short-term incubations (48 h) were carried out on four different days, with each day randomly assigned to a specific temperature and treatment group (CT26, CT32, H226, H232). These incubations were conducted within the shortest possible timeframe lasting from 07^th^ to 23^rd^ of March 2023 to minimize the impact of potential seasonal changes. For each incubation, the respective coral fragments were collected on the same day. Reef environmental parameters, including seawater temperature and salinity were recorded during this period (S1 File) using a multiparameter CTD (Ocean Seven 310, Idronaut, Italy).

### Experimental design

After collection, the coral fragments were transported to the laboratory in transparent sampling bags within a plastic box containing ambient seawater and a portable aquarium pump to ensure aeration. Immediately after arrival, all 24 fragments were attached to custom-made stands marked with colored stripes for the respective colony and placed inside two plastic aquaria, one for each species, filled with 30 L of seawater collected from the reef. The aquaria were equipped with an aquarium pump and a heater connected to a thermostat while natural lighting was simulated with a LED lamp replicating a 12 h light/dark-cycle (PAR ∼ 200 μmol m^-2^ s^-1^, salinity 39.0 ± 1.0 ‰, temperature ambient 26.0 ± 0.5 °C / increased 32 ± 0.5 °C). On each sampling day, all fragments were incubated for 48 h in one treatment (CT = control, H_2_ = hydrogen-enriched) and under one temperature condition (ambient = 26 °C, heat stress = 32 °C) resulting in four different groups: CT26, CT32, H_2_26, H_2_32.

To increase the hydrogen concentration, a hydrogen water generator was used (Hydrogen-rich Water Cup, ABS-FQ-02, Aukewel) that produced 240 mL of hydrogen-enriched distilled water with a concentration of ∼ 3 ppm each cycle according to the manufacturer. In all groups, 10 % of the total seawater was exchanged at the start and after 24 h with artificial seawater created with MilliQ-water and marine sea salt (Red Sea salt, Red Sea) at the respective temperature. In the hydrogen treatments, the artificial seawater was previously hydrogen enriched resulting in a theoretical hydrogen concentration of ∼ 0.3 ppm (∼ 150 µM). After the first water exchange, the temperature was ramped in the heat stress groups from 26 °C to 32 °C within 2 h. Temperature and salinity were checked consistently throughout the experiment using a handheld sensor (YSI professional plus handheld multiparameter meter) to potentially adjust temperature and salinity. Due to a malfunctioning heater in the *P. verrucosa* aquarium during the CT32 group, the water temperature decreased in the first 12 h after the ramping to 28 °C and was then ramped up back to 32 °C within 2 h.

Following the 48 h incubation period, several response parameters related to coral physiology and the overall coral holobiont health were assessed as proxies of the stress state. Initially survival and bleaching of every fragment were visually analyzed. Afterwards six fragments per species were used alive to measure photosynthesis and respiration rates, photosynthetic efficiency (*Fv/Fm*) and rapid light curves (RLC). The remaining six fragments per species were frozen to measure the Symbiodiniaceae cell density and chlorophyll a and c2 content. To normalize response parameters to the size of the coral fragments, the surface area was measured using 3D models in Autodesk ReCap Photo (version 23.1.0) according to Lavy et al. [32] and Tilstra et al. [33]. For this purpose, at least 20 photographs were taken from all sides of the fragment. Photographs were taken after the incubation to avoid stressing the corals beforehand. From these photographs 3D models were created and sliced to only include the fragment surface. The surface area in cm^2^ was then calculated by the program using a reference scale placed on each picture.

### Data collection

#### Survival and bleaching

Survival and bleaching of all fragments were visually assessed directly after the 48 h incubations. Similar to Casey et al. [34], fragments were classified as dead if more than 75 % tissue loss was visible.

#### Photosynthesis and respiration

Six fragments per species were placed on custom-made stands into 1 L jars filled with the respective treatment water and sealed airtight without any air bubbles inside to measure oxygen fluxes according to Tilstra et al. [33] and Mezger et al. [35]. In addition, three 1 L jars per aquaria were filled with treatment water excluding any coral fragments to account for background photosynthesis and respiration rates. Briefly, all jars were placed into a water bath equipped with a heater connected to a thermostat to maintain the temperature at the respective temperature condition. Magnetic stirrers (∼ 200 rpm) placed onto stirring plates (magnetic stirrer HI 200M, Hanna instruments) ensured a constant water flow and homogenous conditions inside the jars. Initially, all fragments and seawater blanks were light-incubated for 90 min (PAR ∼ 200 μmol m^-2^ s^-1^) and afterwards dark-incubated for 60 min. The variation in light and dark incubation period ensured that the oxygen concentration remained within a reliable range. Oxygen concentrations were measured at the start and end of the light and dark incubation using an oxygen probe (Multi 3500i handheld multimeter, WTW and YSI professional plus handheld multiparameter meter). Two calibrated oxygen probes were used in between treatments due to technical difficulties (Multi 3500i handheld multimeter, WTW was used for the H_2_26 group and YSI professional plus handheld multiparameter meter for the CT26, CT32 and H_2_32 groups).

To calculate net photosynthesis (P_net_) and respiration (R) rates, start and end oxygen concentrations of the light and dark incubations were subtracted, respectively. P_net_ and R were then normalized by the surface area of each fragment, total volume of the incubation jar, total incubation time and the background photosynthesis and respiration rates.

Using P_net_ and R, the gross photosynthesis (P_gross_) was then calculated according to following formula:

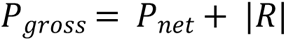

#### Photosynthetic efficiency and rapid light curves

Photosynthetic efficiency was measured directly after the dark incubation using an Imaging-PAM (model IMAG-K7, Walz GmbH) according to Ralph and Gademann [36] and Ralph et al. [37], as the corals were already dark-adapted for >30 min. For each fragment, the dark level fluorescence yield (*F0*) and the maximum fluorescence yield (*Fm*) were measured using the following PAM settings: gain = 2, damp = 2, saturation intensity = 7, saturation width = 4 and measuring light intensity = 7. The optimal PSII quantum yield (*Fv/Fm*) was calculated for five points randomly placed on the coral fragment. In addition, a rapid light curve (RLC) was generated afterwards for the same five points through an exposure of twelve light-series with increasing irradiance from 0 to 702 μmol photons m^−2^ s^−1^. Between the twelve light-series, the fragments were illuminated with the respective actinic light for 20 s, each ended by a saturation pulse. Through all of the five RLCs per fragment one model was fitted according to Platt et al. [38] using the following equation:

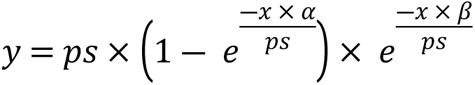

For the model, values of 0 µmol electrons m^-2^ s^-1^ which were recorded at higher irradiances than 0 μmol photons m^−2^ s^−1^ were excluded in advance. This exclusion was made due to the model’s sensitivity as the variables of interest were chosen prior to this range. To determine the variables ps, α and β, a pseudo-random search algorithm was run within the package *phytotools v.1.0* [39] in the program R *v.4.3.0* [40]. The maximum electron transport rate (ETR_max_) and minimum saturating irradiance (E_k_) were calculated using the determined variables as follows:

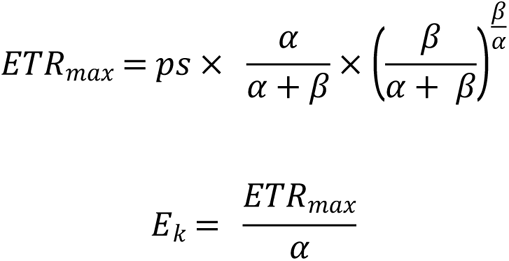

#### Symbiodiniaceae cell densities and chlorophyll concentrations

Coral tissue was removed using an airbrush filled with 10 mL distilled water. The tissue slurry was collected in a 15 mL falcon tube to measure the total volume. In between fragments, the airbrush was cleaned with 70 % ethanol. After airbrushing, the slurries were frozen at −20 °C until further analysis.

To assess the Symbiodiniaceae cell density, a total of 100 µL tissue slurry was thawed and vortexed (analog vortex mixer, VWR). Each sample was then centrifuged for 5 min at 8000 rpm (centrifuge 5424R, Eppendorf). Subsequently, the supernatant was discarded and 1 mL of MilliQ-water was added to the pellet. After dissolving the pellet and homogenizing the mixture using a vortex (analog vortex mixer, VWR), the sample was pipetted on a cell strainer with a 30 µm mesh size to remove larger cells. Following filtration, 200 µL per sample were placed in duplicates in a microplate well. Symbiodiniaceae cell density was then measured using a flow cytometer (Accuri C6 Flow Cytometer, BD). Between samples, a washing and agitating cycle was run and in between three samples, the plate was vortexed to ensure a homogenous mixture. Each sample was run for 2 min and if the events/s recorded by the flow cytometer exceeded 1000, the sample was diluted with MilliQ-water and rerun. The initial count was normalized to the volume sampled by the flow cytometer, surface area, dilution and slurry volume.

A total of 1 mL tissue slurry was thawed and used to measure chlorophyll a and c2 in duplicates. Each sample was centrifuged (centrifuge 5424R, Eppendorf) for 5 min with 5000 rpm at 4 °C and afterwards the supernatant was discarded. To extract chlorophyll a and c2, 2 mL of 90 % acetone was added to each sample. The samples were then sonicated for 10 min and dark-incubated for 24 h at 4 °C. Following 24 h, 200 µL of each sample were transferred in duplicates in a microplate well. Additionally, three 200 µL 90 % acetone blanks were placed on each microplate well to normalize values afterwards. The chlorophyll a and c2 concentrations were assessed by measuring the absorbance at 630 nm (E_630_) and 663 nm (E_663_) using a spectrophotometer (Multiskan SkyHigh microplate spectrophotometer v.1.0.70.0, Thermo Fisher Scientific). Final values were calculated according to Jeffrey and Humphrey [41] and normalized to the surface area, the slurry volume and the averaged absorbance of the blanks. As Jeffrey and Humphrey [41] assumed a path length of 1 cm path which is inaccurate for a microplate well, all values were additionally divided by 0.555. Final chlorophyll a and c2 concentrations were also normalized by Symbiodiniaceae cell density.

#### Statistical analyses

The statistical analyses were conducted in the PRIMER-E software *v.6.1.18* [42] with the PERMANOVA + add on *v.1.0.8* [43] using a permutational multivariate analysis of variance (PERMANOVA). The type III (partial) PERMANOVAs with unrestricted permutations of raw data (999 permutations) including a Monte Carlo test were performed on a resemblance matrix created with the Euclidean distance of the previously normalized data. Two-factorial (factors: temperature [T; two levels: 26 °C, 32 °C], treatment [Tr; two levels: Control, Hydrogen]) PERMANOVAs were carried out for each species on the variables within the oxygen fluxes (P_net_, R, P_gross_), photosynthetic efficiency (*Fv/Fm*), rapid light curves (α, ps/ETR_max_, E_k_), chlorophyll (Chl_a_, Chl_c2_) and Symbiodiniaceae cell density (S1 File). The results were considered significant below a p-value of 0.05. If the interaction effect (T x Tr) was significant, PERMANOVA pairwise tests including a Monte Carlo test were conducted within the temperature and treatment level.

Based on the statistical outcomes, the results were categorized into three groups: the effect of molecular hydrogen (significant main effect of treatment or interaction effect with differences between CT26 and H_2_26), the effect of heat stress (significant main effect of temperature or interaction effect with differences between CT26 and CT32), and the combined effect of molecular hydrogen and heat stress (significant interaction effect with differences between CT32 and H_2_32 or H_2_26 and H_2_32). This orthogonal design allowed for the independent assessment of each factor’s effect and its interaction.

The RLC models and all graphs were created in R *v.4.3.0* [40] using the packages *phytotools v.1.0* [39], *tidyverse v.2.0.0* [44], *ggpubr v.0.6.0* [45] and *rstatix v.0.7.2* [46]. All plots and values represent the mean value and the respective standard error (MEAN ± SE).

## Results

### Effect of molecular hydrogen

Both species, *Acropora* spp. and *P. verrucosa* were affected by hydrogen-enriched seawater at ambient seawater temperatures. In *Acropora* spp., hydrogen-enriched seawater affected the RLCs causing a significant decrease in the ETR_max_ (Fig. 3B, pairwise PERMANOVA, p = 0.001) and E_k_ (Fig. 3C, pairwise PERMANOVA, p = 0.008).

Although the RLCs of *P. verrucosa* remained unaffected by the hydrogen-enriched seawater, similar effects on the photophysiological parameters were observed resulting in a significant decline in the respiration (Fig. 1B, pairwise PERMANOVA, p = 0.001), net photosynthesis (Fig. 1B, pairwise PERMANOVA, p = 0.013) and gross photosynthesis rates (Fig. 1B, pairwise PERMANOVA, p = 0.003) as well as in the areal chlorophyll c_2_ concentration (Fig. 5D, PERMANOVA, p = 0.039).

**Fig. 1.**
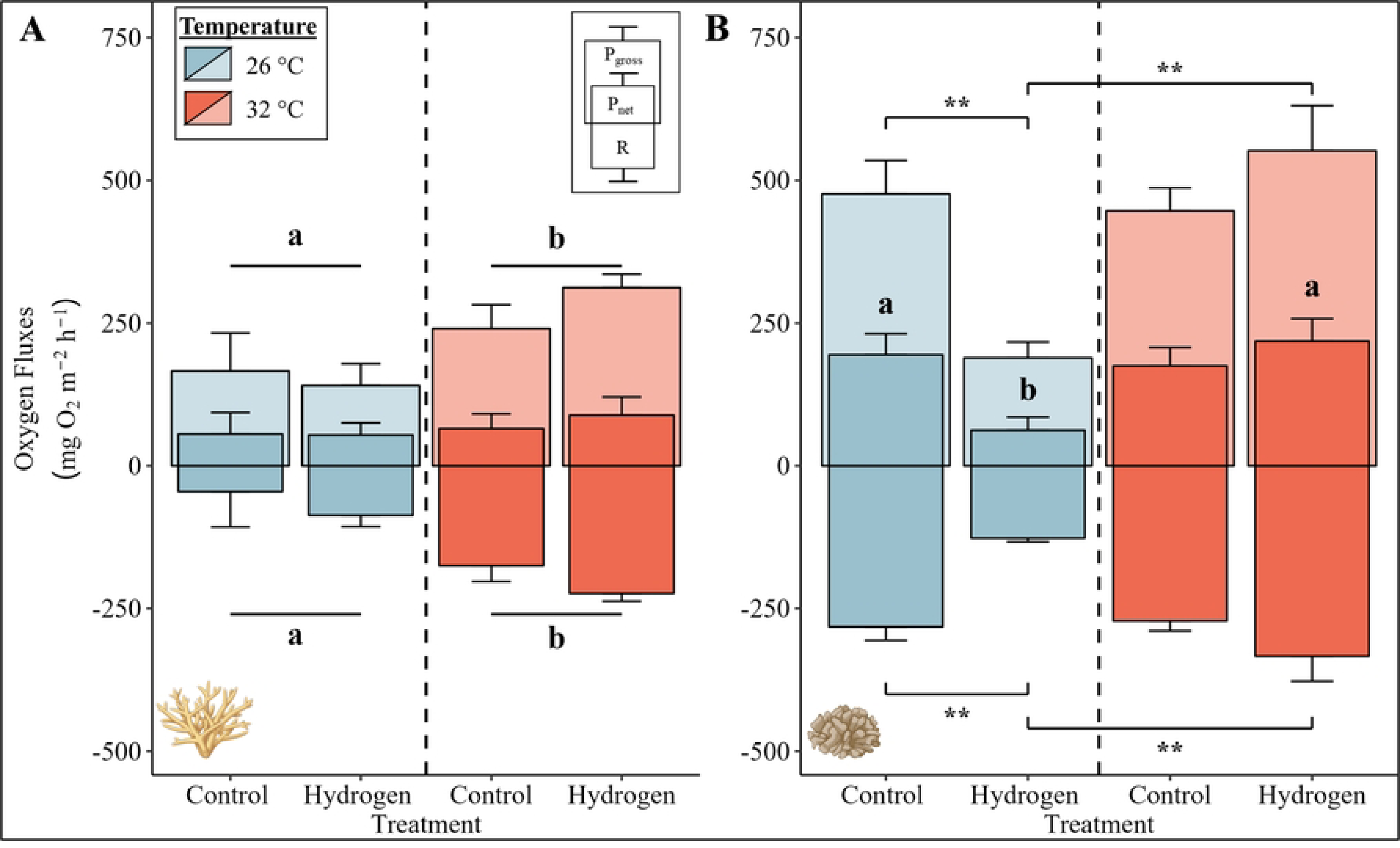
Photosynthesis and respiration rates of (A) *Acropora* spp. and (B) *P. verrucosa* for the four groups. Mean net and gross photosynthesis (P_net_ and P_gross_) and respiration (R) rates are visualized by bars with error bars indicating the respective standard error (SE) of six replica per group (except for five in the CT32 group *P. verruocsa*). The color of the bars indicates the temperature condition (blue = 26 °C, red = 32 °C). A significant temperature main effect according to the PERMANOVA with Monte-Carlo test (p < 0.05) is presented with letters (above bar: P_gross_, below bar: R) and black lines grouping the temperature conditions. Pairwise PERMANOVAs with Monte-Carlo tests were conducted if the interaction effect between temperature and treatment was significant and are illustrated with asterisks (above bar: P_gross_, below bar: R) and letters (P_net_) indicating a significant difference (^a/b,^ * p < 0.05, ** p < 0.01). Exact p-values are presented in supplementary material (S1 File). Coral illustrations were created with BioRender.com.

### Effect of heat stress

While neither species exhibited visual signs of mortality or bleaching across all treatments, heat stress had a significant effect on several physiological response parameters of *Acropora* spp. and *P. verrucosa*. Under heat stress, respiration and gross photosynthesis rates of *Acropora* spp. were significantly increased in both control and hydrogen treatments (Fig. 1A, PERMANOVA, p (respiration) = 0.001, p (gross photosynthesis) = 0.015). In contrast, significantly reduced photosynthetic efficiencies (Fig. 2A, PERMANOVA, p = 0.003), ETR_max_ (Fig. 3B, pairwise PERMANOVA, p = 0.003) and Symbiodiniaceae cell densities (Fig. 5A, PERMANOVA, p = 0.047) were observed in *Acropora* spp. under heat stress in both control and hydrogen treatments. However, chlorophyll a and c2 concentrations remained unaffected (Fig. 5C and 5E).

**Fig. 2.**
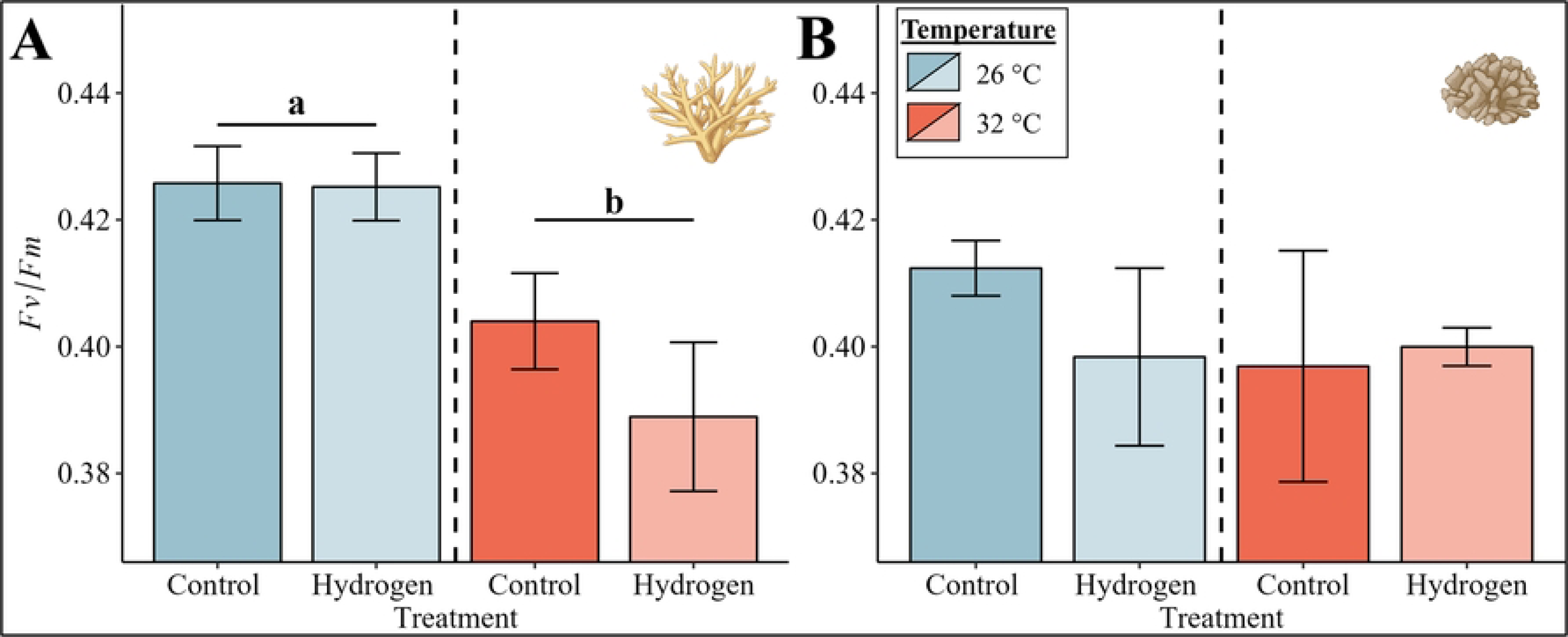
Photosynthetic efficiency (*Fv/Fm*) of (A) *Acropora* spp. and (B) *P. verrucosa* for the four groups. Mean photosynthetic efficiency is visualized by bars with error bars indicating the respective standard error (SE) of six replica per group. The color of the bars indicates the temperature condition (blue = 26 °C, red = 32 °C). A significant temperature main effect according to the PERMANOVA with Monte-Carlo test (p < 0.05) is presented with letters and black lines grouping the temperature conditions. Exact p-values are presented in supplementary material (S1 File). Coral illustrations were created with BioRender.com.

In comparison to the pronounced effect of heat stress in *Acropora* spp., heat stress had a limited effect on *P. verrucosa* as E_k_ was the only parameter to exhibit a significant decrease in both control and hydrogen treatments (Fig. 4B, PERMANOVA, p = 0.018).

### Combined effect of molecular hydrogen and heat stress

Similar to heat stress, the increased temperature caused elevated respiration and gross photosynthesis rates in the combined factor treatment for *Acropora* spp. (Fig. 1A, PERMANOVA, p (respiration) = 0.001, p (gross photosynthesis) = 0.015). Simultaneously, a significant decline in the photosynthetic efficiencies (Fig. 2A, PERMANOVA, p = 0.003) and Symbiodiniaceae cell densities (Fig. 5A, PERMANOVA, p = 0.047) of *Acropora* spp. was observed caused by heat stress. However, in contrast to heat stress and hydrogen addition in which a significant decrease in ETR_max_ of *Acropora* spp. occurred, ETR_max_ remained significantly higher in the combined treatment than in the heat stress (Fig. 3B, pairwise PERMANOVA, p = 0.044) or hydrogen-enriched treatments (Fig. 3B, pairwise PERMANOVA, p = 0.036) and comparable to the rates observed in the ambient control group.

**Fig. 3.**
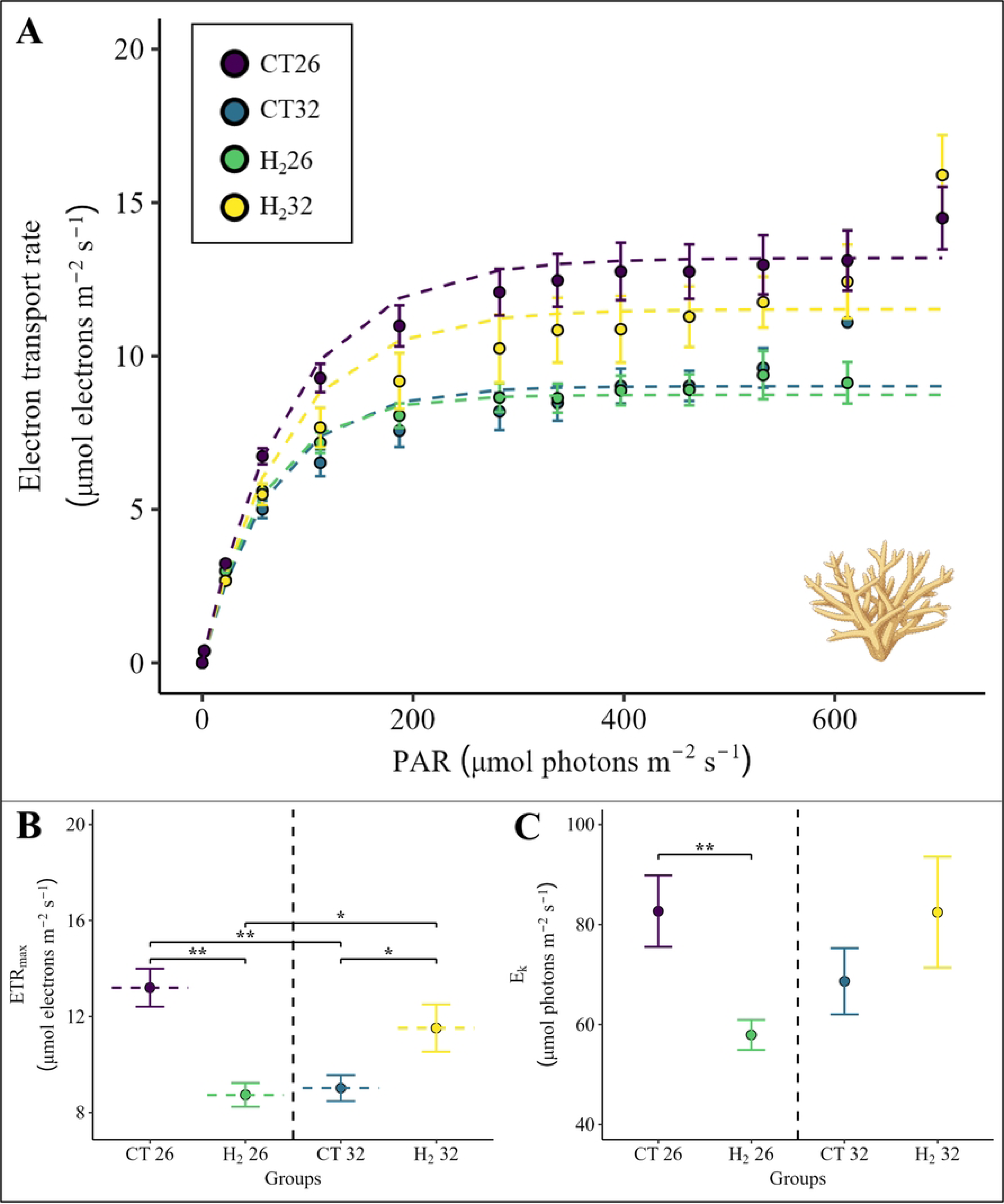
(A) Rapid light curves, (B) maximum electron transport rates (ETR_max_) and (C) minimum saturating irradiances (E_k_) derived from the model fitted according to Platt et al. [38] for all four groups belonging to *Acropora* spp.. The rapid light curves and the respective ETR_max_ and E_k_ values are calculated as the mean of six replica per group (S1 File). The four groups are illustrated with different colors. Points indicate the mean value and error bars the respective standard error (SE). Pairwise PERMANOVAs with Monte-Carlo tests were conducted if the interaction effect between temperature and treatment was significant and are illustrated with asterisks indicating a significant difference (* p < 0.05, ** p < 0.01). Exact p-values are presented in supplementary material (S1 File). Coral illustration was created with BioRender.com.

In the combined treatment of *P. verrucosa*, heat stress was responsible for the significant decline in E_k_ (Fig. 4C, PERMANOVA, p = 0.018) similar to fragments exposed to heat stress alone. Areal chlorophyll c_2_ concentrations were also significantly reduced in the combined treatment of *P. verrucosa* (Fig. 5D, PERMANOVA, p = 0.039) although not attributed to heat stress but to hydrogen addition similar to fragments exposed to hydrogen addition alone. However, respiration, net photosynthesis and gross photosynthesis were affected by an interactive effect of treatment and heat stress (Fig. 1B, PERMANOVA, p (respiration) = 0.001, p (net photosynthesis) = 0.019, p (gross photosynthesis) = 0.003). Although the addition of hydrogen alone caused a significant decline, the combined effects of treatment and heat stress resulted in notably higher respiration, net photosynthesis and gross photosynthesis rates (Fig. 1B, pairwise PERMANOVA, p (respiration) = 0.001, p (net photosynthesis) = 0.011, p (gross photosynthesis) = 0.001) comparable to those observed in the ambient and heat stress control groups.

**Fig. 4.**
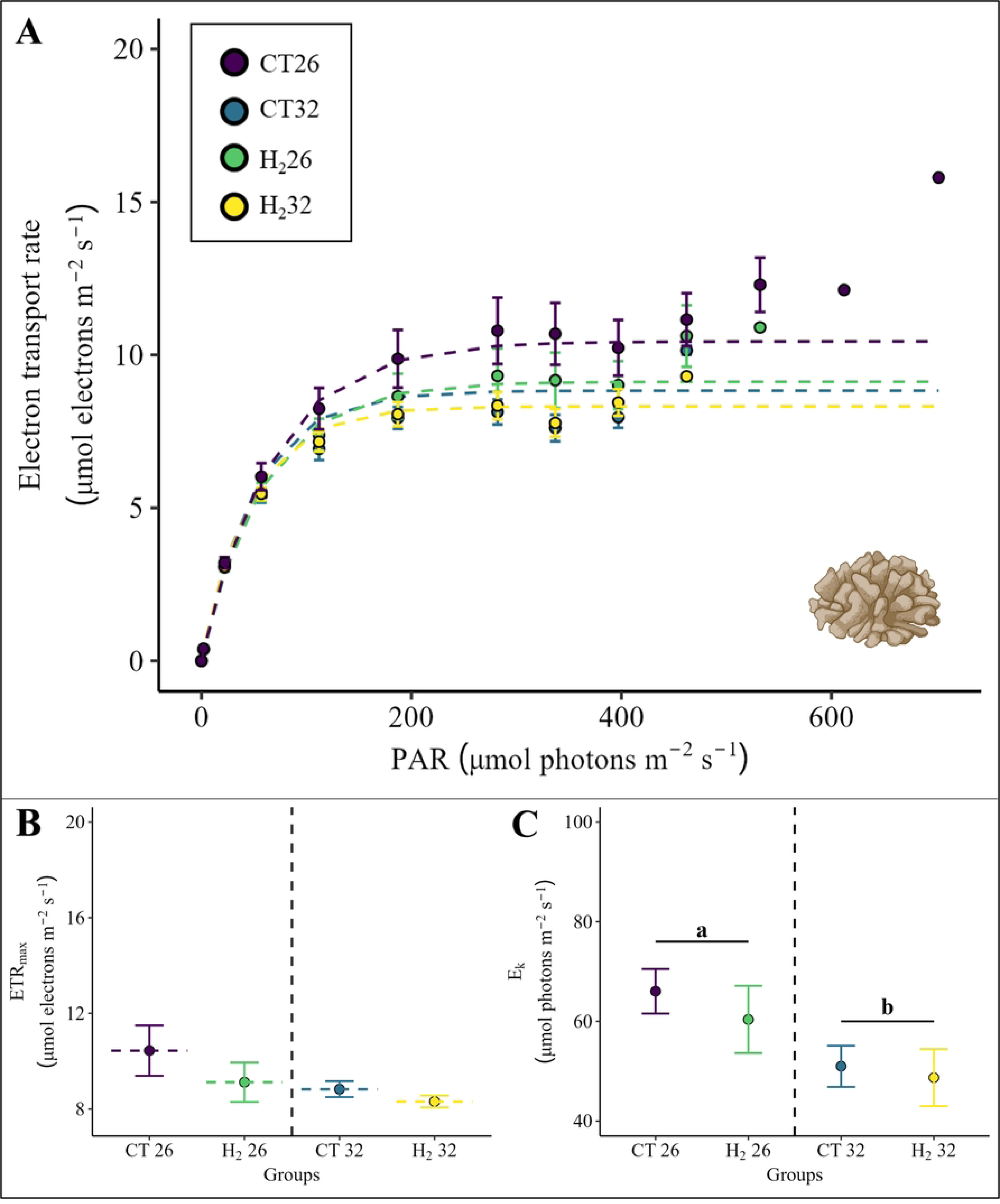
(A) Rapid light curves, (B) maximum electron transport rates (ETR_max_) and (C) minimum saturating irradiances (E_k_) derived from the model fitted according to Platt et al. [38] for all four groups belonging to *P. verrucosa*. The rapid light curves and the respective ETR_max_ and E_k_ values are calculated as the mean of six replica per group (S1 File). The four groups are illustrated with different colors. Points indicate the mean value and error bars the respective standard error (SE). A significant temperature main effect according to the PERMANOVA with Monte-Carlo test (p < 0.05) is presented with letters and black lines grouping the temperature conditions. Exact p-values are presented in supplementary material (S1 File). Coral illustration was created with BioRender.com.

**Fig. 5.**
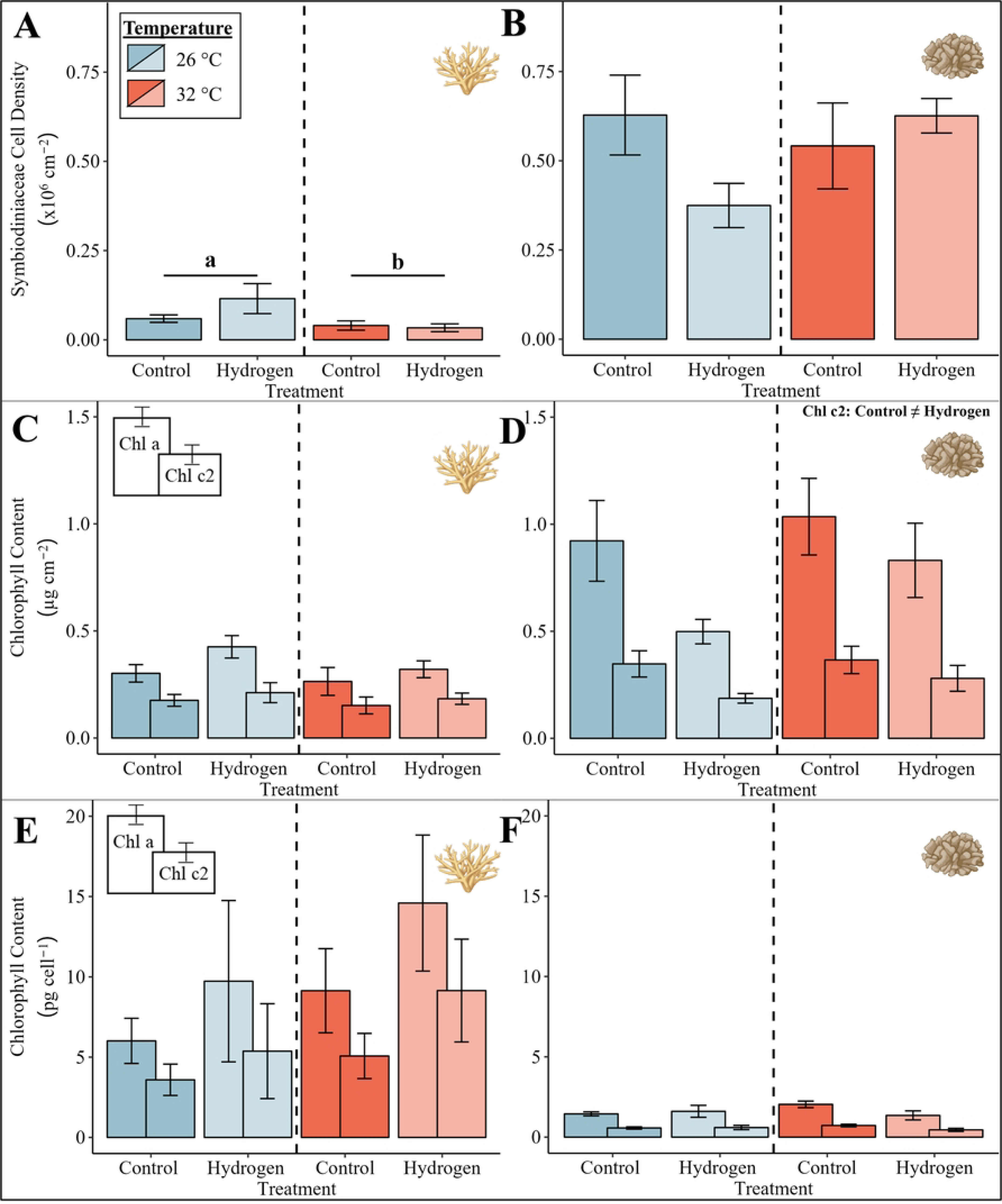
Symbiodiniaceae cell density of (A) *Acropora* spp. and (B) *P. verrucosa*, areal chlorophyll content of (C) *Acropora* spp. and (D) *P. verrucosa* and chlorophyll content per cell of (E) *Acropora* spp. and (F) *P. verrucosa* for the four groups. Bars with error bars indicate the mean and the respective standard error (SE) of six replica per group. For the chlorophyll content, chlorophyll a is presented on the left side of each bar with chlorophyll c2 at the right side. The color of the bars indicates the temperature condition (blue = 26 °C, red = 32 °C). A significant temperature main effect according to the PERMANOVA with Monte-Carlo test (p < 0.05) is presented with letters and black lines grouping the temperature conditions. A significant treatment main effect according to the PERMANOVA with Monte-Carlo test (p < 0.05) is presented with the unequal sign in the top right corner. Exact p-values are presented in supplementary material (S1 File). Coral illustrations were created with BioRender.com.

## Discussion

As coral reefs currently face the fourth global coral bleaching event [17, 47] reaffirming the catastrophic projections for an increase in both scale and frequency of such events [10, 15, 16, 17], it becomes crucial to investigate new approaches strengthening coral reefs thereby potentially counteracting these events in the future [48, 49]. In this context, molecular hydrogen might offer a promising approach since several studies have already demonstrated an extensive preventive and therapeutic effect of hydrogen in mammals including humans and rats [23, 24, 25, 26, 27, 28]. Our study is therefore the first to investigate the short-term effects of hydrogen on the coral holobiont ecophysiology under ambient temperatures and heat stress, thereby providing an overview of the potential ecological implications.

### Molecular hydrogen can minimize negative effects of heat stress on the hard coral *Acropora* spp

When subjected to heat stress alone, *Acropora* spp. experienced a significant reduction in the photosynthetic efficiency, ETR_max_ and Symbiodiniaceae cell density alongside an increase in metabolic demand - indicative of a pronounced stress response [50, 51]. However, simultaneous exposure to hydrogen and heat stress prevented the decline in the ETR_max_ observed with heat stress alone in *Acropora* spp.. As shown in previous studies, a decrease in ETR_max_ can serve as an indicator of damage to the photosystem and thereby a deteriorated health state of the Symbiodiniaceae cells [52, 53]. This damage presumably originates from the accumulation of ROS/RNS under heat stress as proposed by the “Oxidative Theory” [20, 21, 22, 54, 55]. Our result therefore suggests that molecular hydrogen might have protected the coral holobiont against oxidative damage, thereby preventing a decrease in ETR_max_. These findings align with the previously reported antioxidative capacity of molecular hydrogen [23, 24, 25, 26, 27, 28]. In addition, molecular hydrogen may protect the photosynthetic apparatus indirectly by increasing antioxidant enzyme activity [56, 57] and upregulating the heat shock response as observed in plants and rodents, respectively [31].

Consistent with our observations, a previous study in the Central Red Sea found that *Acropora* spp. was more affected by the simulated short-term heat stress in comparison to *P. verrucosa* [58]. This different response of *P. verrucosa* could indicate that the simulated heat stress was insufficient to induce negative effects within 48 h and suggests a higher temperature resistance in *P. verrucosa* even though other studies have described *P. verrucosa* as more sensitive to temperature stress than *Acropora hemprichii* in the Red Sea [59].

### Molecular hydrogen negatively affects the photophysiology of *Acropora* spp. and *P. verrucosa* under ambient seawater temperatures

Interestingly, molecular hydrogen also affected the photophysiology of *Acropora* spp. and *P. verrucosa* under ambient seawater temperatures. In several *in vitro* studies, molecular hydrogen was observed to inhibit nitrogen fixation in diazotrophs by competing with atmospheric nitrogen thereby decreasing the formation of biologically active nitrogen in the form of ammonium [60, 61, 62, 63]. This process, occurring in diazotrophs within the coral holobiont, is an important process to supply bioavailable nitrogen [64, 65] and therefore hydrogen’s potential to inhibit nitrogen fixation rates might have limited the overall nitrogen availability of the coral holobiont under ambient seawater temperatures. As the photophysiology and growth are usually limited by nitrogen availability [66, 67, 68, 69, 70, 71, 72, 73], this might account for the significantly lower photosynthesis rates and chlorophyll c2 concentrations in *P. verrucosa* and a significant decline in symbiont photophysiology of *Acropora* spp. when exposed to hydrogen. Lower respiration rates of *P. verrucosa* might directly correlate to the reduced primary productivity as less photosynthates are available for respiration [74]. Overall, molecular hydrogen had a more pronounced effect on *P. verrucosa* under ambient temperatures which might relate to the finding that nitrogen fixation rates were ∼ 3-fold higher in *Acropora* spp. than in *P. verrucosa* [75] and therefore a reduction of nitrogen fixation rates through the addition of hydrogen may have a greater impact on the already lower nitrogen fixation rates of *P. verrucosa*.

Under the combined treatment of hydrogen addition and heat stress, the photosynthesis and respiration rates in *P. verrucosa* were significantly higher compared to hydrogen addition under ambient seawater temperatures and similar to the rates observed in the ambient and heat stress control treatments. At higher temperatures, nitrogen fixation rates in the coral holobiont were shown to increase [76, 77, 78] and therefore the potentially reduced nitrogen fixation by molecular hydrogen [60, 61, 62, 63] could have balanced the nitrogen availability resulting in a higher productivity than under hydrogen addition at ambient temperatures. This effect might even be beneficial at heat stress as an increased nitrogen availability cause an uncontrollable proliferation of endosymbionts [51] disrupting the usual nitrogen-limited state within the coral holobiont [19, 78, 79, 80, 81]. Molecular hydrogen might therefore support maintaining the nitrogen limitation essential for the successful symbiosis under heat stress.

In contrast, Rädecker et al. [82] could demonstrate that the ammonium, produced by the increased nitrogen fixation rates under heat stress, was not assimilated by the endosymbionts. Considering these circumstances, the inhibitory effect of hydrogen would not support the endosymbionts under heat stress but potentially maintain the balanced nutrient concentration in the whole coral holobiont as the increased nitrogen availability might influence other groups like nitrifiers or denitrifiers [79, 83].

### Ecological Implications

Overall, these findings indicate a potential negative effect of hydrogen addition under ambient seawater temperatures on the coral holobiont ecophysiology of *Acropora* spp. and *P. verrucosa* presumably due to the inhibition of nitrogen fixation. However, under heat stress hydrogen had a beneficial effect on the symbiont photophysiology of *Acropora* spp., indicating that molecular hydrogen is incorporated and potentially strengthens coral physiology. This demonstrates the potential of molecular hydrogen and paves the way for future long-term studies potentially contributing to the development of new management strategies aiming at enhancing coral resilience to counteract negative effects of heat stress. This raises the question about the safety, cost, potential administration methods and the mechanisms behind molecular hydrogen (Hypothesized mechanisms in Fig. 6).

**Fig. 6.**
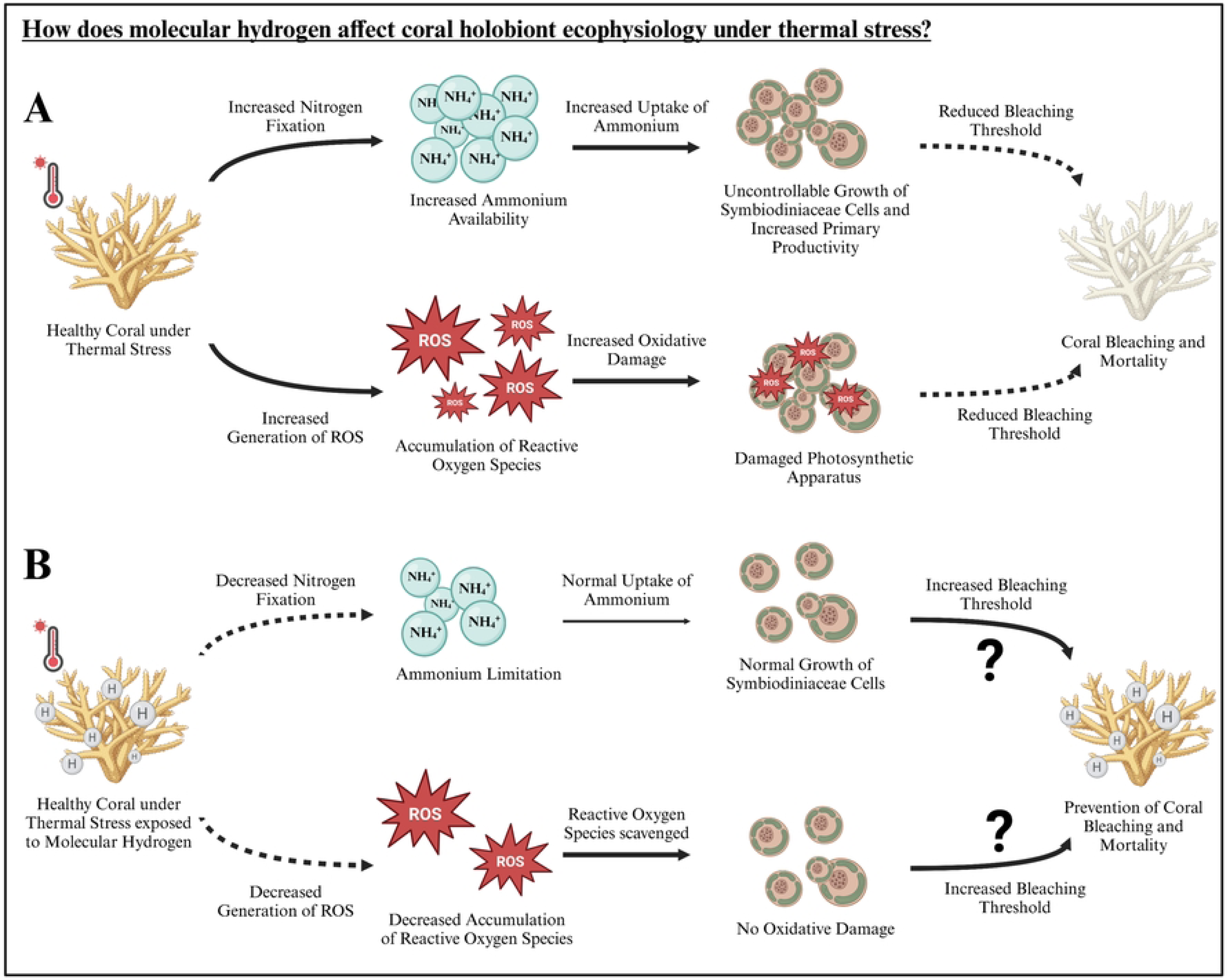
Hypothesized effect of (A) heat stress and (B) heat stress in combination with molecular hydrogen on the coral holobiont ecophysiology. First results indicate that hydrogen prevents oxidative damage of the photosystem of the Symbiodiniaceae cells. The potential mechanism of molecular hydrogen behind this effect is illustrated above with dotted lines indicating a decrease and bold lines indicating an increase in the respective process compared to each other. The question mark indicates a potential long-term effect which needs to be addressed in further studies (Illustration created with BioRender.com).

Most importantly, molecular hydrogen should only be supplied during heat stress events with hydrogen gas concentrations lower than 4 % to minimize the risk of explosion as molecular hydrogen is highly flammable above these concentrations [84]. Due to hydrogen’s short retention time [85], corals should therefore only be affected directly after the administration and potential negative effects at ambient seawater temperatures might be prevented. Since hydrogen is a very small molecule rapidly diffusing across cell membranes [23], it is very likely that hydrogen will also affect other reef organisms. A recent study, for instance, demonstrated that hydrogen promotes the growth of diverse marine bacteria [86]. Therefore, the effects of molecular hydrogen on the microbiome as well as other invertebrates and vertebrates, as well as the underlying mechanisms driving the enhanced thermal resistance, must be clarified prior to application.

Since hydrogen gained a lot of attention in recent years as a potential emission-free and renewable energy carrier [24, 87], considerable research has been conducted to develop simple and affordable methods for hydrogen production. Electrolysis devices converting distilled water or even seawater [88, 89] into molecular hydrogen might be transported to vulnerable reefs bubbling hydrogen directly into the reef. In addition, hydrogen tablets [90] might be placed on particularly threatened corals constantly supplying hydrogen. Another possibility to provide thermal-stressed corals with hydrogen is supplementation via other approaches. Microbiome interventions particularly reshaping the microbiome through probiotics are emerging as very promising tools for coral reef conservation [91, 92, 93, 94]. Given the widespread abundance of hydrogen-producing bacteria [95] and their likely presence within the coral microbiome, identifying these bacteria could reveal new beneficial microorganisms potentially further supporting the development of effective probiotics.

However, before an *in vivo* application is feasible, further research is necessary as this study only demonstrates a short-term effect of molecular hydrogen, and potential negative effects under ambient temperature. First, the long-term effects of molecular hydrogen on the coral holobiont, along with potential effects on other reef organisms, have to be investigated. If successful, the underlying mechanisms, particularly related to oxidative stress and nitrogen availability, must be deciphered in combination with an ideal concentration range of molecular hydrogen to implement a potential *in vivo* application.

## Acknowledgments

We would like to thank the whole Marine Microbiomes working group for their great assistance during this work. A big thanks also goes to the boat crew and scientific diving team of the Red Sea Research Center as well as the Coastal and Marine Resources Core Lab (KAUST) for their support in all marine operations.

## Supporting Information

**S1 File. Supplementary figures and statistical tables.**

**S1 Data.**

## Notes

### Competing Interest Statement

The authors have declared no competing interest.

